# ANTAGONISTIC ROLES FOR ATAXIN-2 STRUCTURED AND DISORDERED DOMAINS IN RNP CONDENSATION

**DOI:** 10.1101/2020.07.02.184796

**Authors:** Amanjot Singh, Joern Huelsmeier, Arvind Reddy Kandi, Sai Shruti Pothapragada, Jens Hillebrand, Arnas Petrauskas, Khushboo Agrawal, RT Krishnan, Devasena Thiagarajan, K. VijayRaghavan, Mani Ramaswami, Baskar Bakthavachalu

## Abstract

Ataxin-2 is a conserved translational control protein associated with spinocerebellar ataxia type II (SCA2) and amyotrophic lateral sclerosis (ALS) as well as an important target for ALS therapeutics under development. Despite its clinical and biological significance, Ataxin-2’s activities, mechanisms and functions are not well understood. While *Drosophila* Ataxin-2 (Atx2) mediates mRNP condensation via a C-terminal intrinsically disordered domain (cIDR), how Ataxin-2 IDRs work with structured (Lsm, Lsm-AD and PAM2) domains to enable positive and negative regulation of target mRNAs remains unclear. Using TRIBE (Targets of RNA-Binding Proteins Identified by Editing) technology, we identified and analysed Atx-2 target mRNAs in the *Drosophila* brain. We show that Atx2 preferentially interacts with AU-rich elements (AREs) in 3’UTRs and plays a broad role in stabilization of identified target mRNAs. Strikingly, Atx2 interaction with its targets is dependent on the cIDR domain required for neuronal-granule formation. In contrast, Atx2 lacking its Lsm domain not only interacts more efficiently with the target mRNA identified, but also forms larger RNP granules. Providing an extensive dataset of Atx2-interacting brain mRNAs, our results demonstrate that Atx2: (a) interacts with target mRNAs within RNP granules; (b) modulates the turnover of these target mRNAs; (c) has an additional essential role outside of mRNP granules; and (d) contains distinct protein domains that drive or oppose RNP-granule assembly. These findings increase understanding of neuronal translational control mechanisms and inform Ataxin-2-based interventions in development for SCA2 and ALS.

## INTRODUCTION

Ataxin-2’s involvement in human disease, its relevance for therapeutics development and its established roles in mRNP-phase transitions, cell physiology, metabolic control and animal behavior, have led to considerable interest in understanding molecular mechanisms by which the protein functions. At a molecular level, Ataxin-2 positively or negatively regulates the translation of specific mRNAs (Lee et al., 2017; Lim and Allada, 2013; McCann et al., 2011; Zhang et al., 2013). At the same time, the protein mediates the assembly of mRNPs into cytoplasmic mRNP granules visible in resting neurons or in RNA stress granules (SGs) that occur in most cells in response to stress (Bakthavachalu, Huelsmeier et al., 2018; Bentmann et al., 2013). At a cellular level, Ataxin-2 contributes to cell viability and differentiation as well as cellular responses to viral, ER, heat- and oxidative stress (Bonenfant et al., 2019; del Castillo et al., 2019; Ralser et al., 2005; van de Loo et al., 2009). Finally, at an organismal level, the protein regulates metabolism, circadian rhythm and the consolidation of long-term memory (Bakthavachalu, Huelsmeier et al., 2018; Lim and Allada, 2013; Meierhofer et al., 2016; Pfeffer et al., 2017; Zhang et al., 2013). Parallel clinical genetic studies have shown that genetic mutations in human Ataxin-2 (Atxn2) can cause the hereditary neurodegenerative diseases spinocerebellar ataxia type II (SCA2) or amyotrophic lateral sclerosis (ALS) (Daoud et al., 2011; Elden et al., 2010; Lastres-becker et al., 2007; Lee et al., 2011; Scoles and Pulst, 2018; Wadia, 1977; Wadia and Swami, 1971), and subsequent work showing that genetic reduction of Ataxin-2 activity slows neurodegeneration in animal models of ALS has inspired the design and development of therapeutics targeting human Ataxin-2 (Becker et al., 2017; Becker and Gitler, 2018; Elden et al., 2010; Scoles et al., 2017).

The above biological and clinical studies of Ataxin-2 are connected by the insight that intrinsically disordered domains (IDRs) present on RNA-binding proteins contribute to macromolecular condensation or liquid-liquid phase separation reactions, wherein monomeric units form dynamic assemblies held together by weak multivalent interactions (Jain and Vale, 2017; Kato and McKnight, 2018; Murray et al., 2017; Saha and Hyman, 2017; Van Treeck et al., 2018; Van Treeck and Parker, 2018). Significantly, IDRs not only support assembly of mRNP granules but also are prone to assemble into amyloid-like fibers. Disease causing mutations often increase the efficiency of amyloid formation, particularly within mRNP granules where the RNA-binding proteins are concentrated (Courchaine et al., 2016; Kato et al., 2012; Lim et al., 2019; Nedelsky and Taylor, 2019; Patel et al., 2015; Ramaswami et al., 2013; Xiang et al., 2015; Yang et al., 2018). The broad proposal that SGs serve as “crucibles” for initiation of neurodegenerative disease (Li et al., 2013; Ramaswami et al., 2013; Wolozin and Ivanov, 2019) explains why SG proteins are often mutated in familial ALS or Frontotemporal Dementia (FTD) and why these proteins are observed in intracellular inclusions typical of ALS/FTD (Arai et al., 1992; Brettschneider et al., 2015; Lee et al., 1991).

The domain structure of Ataxin-2 is highly conserved across species, with N-terminal Lsm (Like-Sm) and LsmAD (Lsm-associated) domains, a more carboxy-terminal PAM2 (polyA binding protein interaction motif 2) domain as well as strongly disordered regions (respectively mIDR and cIDR) in the middle and C-terminal regions of the protein (Albrecht et al., 2004, Satterville and Pallanck, 2006, Nonhoff et al., 2007; Bakthavachalu, Huelsmeier et al., 2018). Genetic studies in *Drosophila*, which has a single gene for Ataxin-2 as against two *atxn2* and *atxn2-like* in mammals, indicate that different Atx2 domains encode distinct, biological functions. Specifically, while each structured domain is essential for normal viability, the IDR domains are not. However, the cIDR is required for normal mRNP assembly and long-term memory as well as for facilitating cytotoxicity in *Drosophila* Fus and C9orf72 models for ALS/FTD (Bakthavachalu, Huelsmeier et al., 2018). These observations, while instructive in terms of functions of the Atx2-IDR and mRNP granules, provide no direct insight into other functions and mechanisms mediated by structured domains of Atx2 or their roles in biology.

To better understand broad roles and mechanisms of Atx2, we used TRIBE (Targets of RNA-Binding Proteins Identified by Editing) technology (Biswas et al., 2019; McMahon et al., 2016) to globally identify Atx2 interacting mRNAs from *Drosophila* adult brain and study how these *in vivo* interactions were influenced by different domains of the protein. In addition to identifying biologically important targets of Ataxin-2, results described here offer unexpected information into mechanisms of Atx2 protein function. Atx2 associates with mRNAs predominantly within mRNP granules, where it binds preferentially near AU rich elements (AREs) in the 3’UTRs to stabilize the majority of the targets. While the cIDR enables mRNA interactions and granule assembly the Lsm domain reduces both mRNP assembly and Atx2-target interactions. Taken together, our data: (a) provide a rich data set of Atx2 target mRNAs; point to a novel essential function of Ataxin-2 outside of mRNP granules; and (c) indicate competing disassembly and pro-assembly activities within Atx2 encoded by the Lsm and IDR domains respectively. In addition to being of specific biological interest, these conclusions are relevant to current therapeutic strategies based on targeting human Atxn2.

## RESULTS

### Using TRIBE to identify Atx2-target mRNAs in Drosophila brain

Atx2 is abundantly expressed in brain tissue. To identify *in vivo* targets of Atx2 in the fly brain, we used TRIBE, a technology previously shown to reproducibly identify RBP target mRNAs *in vivo* (McMahon et al., 2016). We generated transgenic flies that express Atx2 linked to the catalytic domain of *Drosophila* RNA-modifying enzyme, adenosine deaminase (ADARcd) at the carboxy terminal along with a V5 epitope tag under the control of the Gal4-responsive UAS promoter (Suppl. Figure S1A). In tissue expressing the Atx2-ADARcd transgene, mRNAs should undergo Adenosine-to-Inosine editing specifically at positions proximal to Ataxin-2 binding sites (Figure 1A). In an *elav-Gal4* background, where the Gal4 transcription factor is expressed in postmitotic neurons, Atx2-ADARcd is expressed specifically in the nervous system. We further temporally restricted neural expression to adult flies with the use of the temperature sensitive Gal4 inhibitor, GAL80^ts^ that is active at temperatures below 25°C. Thus, in *elav-Gal4; TubGal80^ts^, UAS-Atx2-ADARcd* adult flies shifted from 18°C to 29°C for 5 days shortly after eclosion, neural mRNAs expressed in adult flies would be susceptible to editing at adenosine residues proximal to Atx2 binding sites.

**Figure 1:**
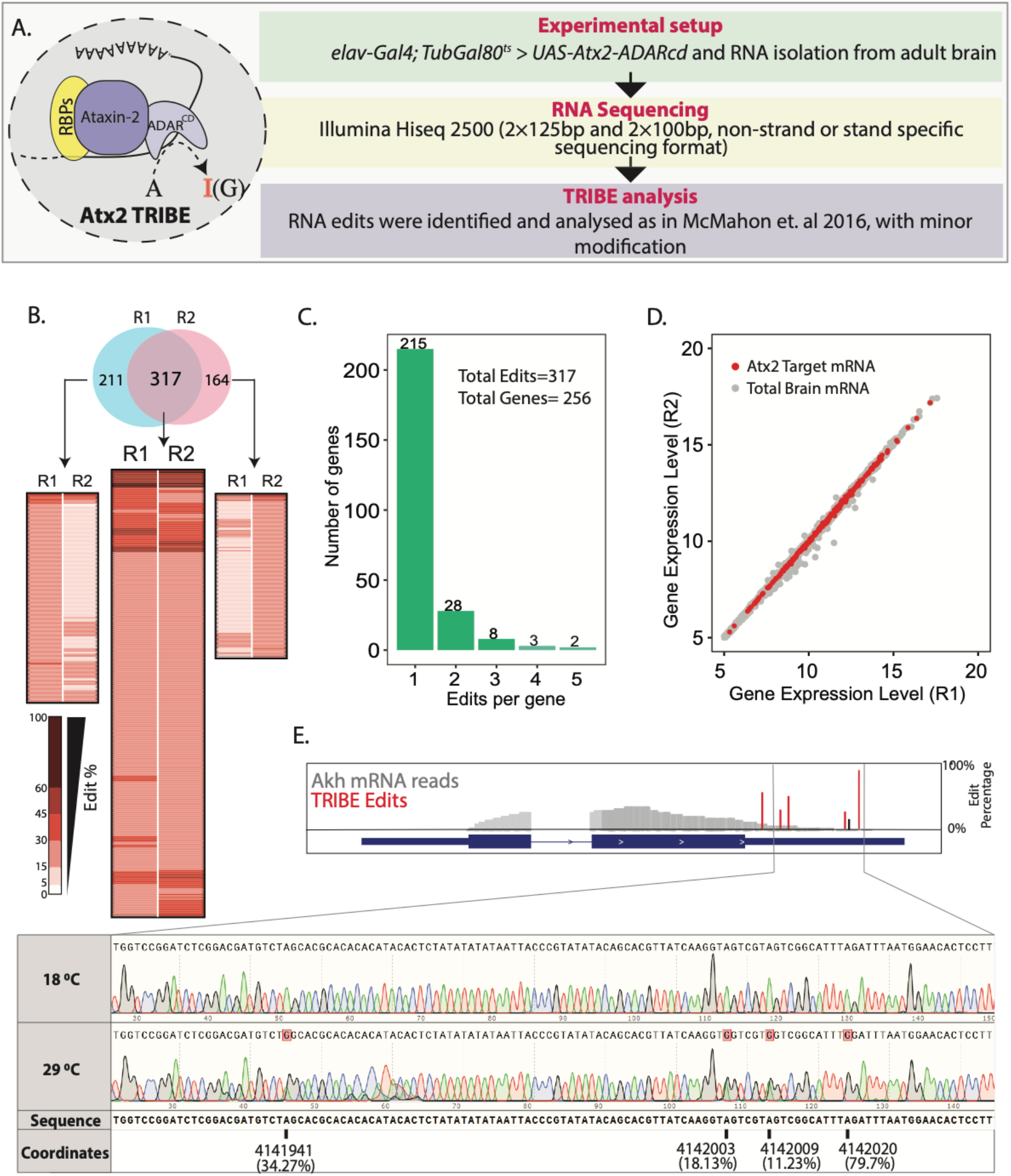
Using TRIBE to identify Atx2-interacting mRNAs in the adult fly brain. (A) Schematic and flow chart for TRIBE analysis: the Atx2-ADARcd fusion protein is expressed in the adult fly brain, total brain mRNA isolated, sequenced and analysed using a published TRIBE pipeline. (B) Heatmaps show edit percentages of individual transcript coordinates. Replicate experiments R1 and R2 identify largely overlapping edit sites and edited mRNAs. The common targets (intersect between R1 and R2) show almost reproducible edit levels. The inset (heat maps on either sides of the common intersected list) indicates that several mRNAs are identified as “non-replicates” between R1 or R2, because they do not cross quality-control thresholds (edit percentages or read-counts) and not because of robust differences between replicates (C) Bar plot showing the number of edits with a significant number of genes edited at a single site. (D) The mRNAs edits are independent of expression levels. A scatter plot shows expression differences of all the mRNA expressed in fly brain. Red dots represent the edited mRNAs while grey dots represent mRNAs of the brain transcriptome which are not edited. (E) Sanger sequencing confirms editing site identified by TRIBE analysis. Data shown for one target mRNA (Akh). Red bars show the edit percentages at the different modified nucleotides with respect to the total Akh mRNA shown in grey. The black bar indicates identified edits that are below 15% threshold (See also Suppl. Figure S1D).

To identify neural mRNA targets of Atx2, we isolated polyA-selected RNAs from Atx2-ADARcd expressing *Drosophila* brain and sequenced these using Illumina Hiseq 2500 and reads were subsequently analyzed according to McMahon et al 2016 (with slight modifications, see suppl. methods) to identify sites and efficiency of Atx2-ADARcd mediated mRNA editing (Figure 1A). Adult brain expression for Atx2-ADARcd was verified using antibody staining for V5 epitope (Supp Figure 1B). Experiments were carried out in duplicates. The reads obtained were between 20 and 25 million per sample, were of high quality and more than 80% of them mapped to specific *Drosophila melanogaster* mRNAs (Supp Figure 1C).

Transcripts from about 8% of expressed genes showed adenosine edits. These edits required expression of the full Atx2-ADARcd fusion protein and were not consequences of background, endogenous or non-specific ADAR activity. Of the 528 and 481 edit sites represented in independent replicates, at least 317 were common to both samples, demonstrating the reproducibility of the experiments (Figure 1B). These 317 common edits could be assigned to 256 unique genes, with the majority of genes only being edited at a single site (215 genes, 67.8% of edits). The Atx2 target mRNAs were distributed across the brain transcriptome irrespective of the abundance of a mRNA indicating that edits were not random events, but rather reflect sequence and structure specific association of these mRNAs with Atx2-ADARcd (Figure 1D). We further validated the TRIBE analysis/ Illumina Hiseq pipeline using Sanger sequencing for a few identified target mRNAs. For instance, Sanger sequencing confirmed that edits in *Adipokinetic hormone* (*Akh*) and *14-3-3 epsilon* mRNAs occur at identified sites, with efficiencies indicated by TRIBE analysis (Figure 1E; Suppl. Figure 1D). Taken together, the data indicates that Atx2 associates with specific RNA motifs present on targets identified by TRIBE analyses from fly brain.

A gene ontology analysis of Atx2-targets (Table 1) indicated particular enrichment of mRNAs encoding neuropeptides and hormones, as well as mRNAs encoding monoamine transporters, ion channels and vesicle transport proteins. This is broadly consistent with and suggests mechanisms by which Atx2 functions in translational control of physiological and neural circuit plasticity.

**Table 1:** Gene Ontology analysis of Atx2 brain targets. GO analysis using PANTHER shows a large enrichment of neuronal mRNAs coding for neuropeptides and proteins involved in neuronal signaling pathways. (FDR = False Discovery Rate).

| Molecular Function | Fold enrichment | Probability (FDR) | Targets (%) | Genome (%) | GO Term |
| --- | --- | --- | --- | --- | --- |
| signaling receptor binding | 7.75 | 4.47E-11 | 14.46 | 1.87 | GO:0005102 |
| neuropeptide hormone activity | 27.67 | 2.52E-09 | 6.63 | 0.24 | GO:0005184 |
| hormone activity | 18.64 | 5.74E-08 | 6.63 | 0.36 | GO:0005179 |
| neuropeptide receptor binding | 38.74 | 2.08E-06 | 4.22 | 0.11 | GO:0071855 |
| G protein-coupled receptor binding | 13.84 | 3.79E-06 | 6.03 | 0.44 | GO:0001664 |
| receptor regulator activity | 9.58 | 4.55E-06 | 7.23 | 0.76 | GO:0030545 |
| receptor ligand activity | 9.41 | 1.96E-05 | 6.63 | 0.71 | GO:0048018 |
| signaling receptor activator activity | 9.04 | 2.58E-05 | 6.63 | 0.74 | GO:0030546 |
| organic hydroxy compound transmembrane transporter activity | 83.02 | 3.38E-04 | 2.41 | 0.03 | GO:1901618 |
| monoamine transmembrane transporter activity | 66.41 | 5.52E-04 | 2.41 | 0.04 | GO:0008504 |
| phosphoric ester hydrolase activity | 4.93 | 6.43E-04 | 7.84 | 1.59 | GO:0042578 |
| anion:sodium symporter activity | 55.35 | 6.44E-04 | 2.41 | 0.05 | GO:0015373 |
| sodium:chloride symporter activity | 55.35 | 6.84E-04 | 2.41 | 0.05 | GO:0015378 |
| phosphoprotein phosphatase activity | 7.95 | 7.06E-04 | 5.43 | 0.69 | GO:0004721 |
| anion:cation symporter activity | 36.9 | 1.81E-03 | 2.41 | 0.07 | GO:0015296 |
| cation:chloride symporter activity | 36.9 | 1.91E-03 | 2.41 | 0.07 | GO:0015377 |
| phosphatase activity | 5.05 | 2.53E-03 | 6.63 | 1.32 | GO:0016791 |
| serotonin:sodium symporter activity | 83.02 | 4.17E-03 | 1.81 | 0.03 | GO:0005335 |
| neurotransmitter: sodium symporter activity | 14.82 | 5.04E-03 | 3.02 | 0.21 | GO:0005328 |
| protein tyrosine/serine/threonine phosphatase activity | 23.72 | 6.17E-03 | 2.41 | 0.11 | GO:0008138 |
| solute: sodium symporter activity | 11.53 | 1.26E-02 | 3.02 | 0.27 | GO:0015370 |
| mRNA 3'-UTR binding | 11.22 | 1.36E-02 | 3.02 | 0.27 | GO:0003730 |
| protein tyrosine phosphatase activity | 10.64 | 1.59E-02 | 3.02 | 0.29 | GO:0004725 |
| neurotransmitter transmembrane transporter activity | 10.64 | 1.65E-02 | 3.02 | 0.29 | GO:0005326 |
| adrenergic receptor activity | 24.91 | 3.98E-02 | 1.81 | 0.08 | GO:0004935 |

### Atx2 associates preferentially with 3’UTRs of target mRNAs

The RNA edit sites identified by TRIBE reflect positions to which ADARcd is targeted via direct or indirect Atx2-mRNA interactions. It is therefore possible to use this information to determine the relative positions and preferred sequences for Atx2 binding on respective mRNAs. By converting the TRIBE edits into metagene coordinates, we found that Atx2 interactions occurred predominantly with 3’UTRs of target mRNAs (69.5%), while the coding region (CDS) and 5’UTR accounted for 26.7% and 4% of edits respectively (Figure 2A). All the identified edits occurred almost exclusively in exons (Suppl. Figure 2A) suggesting that Atx2 binds to mature mRNA in the cytoplasm. Edits within CDS were often accompanied by edits in the 3’UTR of the same mRNA, further indicating a key role for Atx2-3’UTR interactions (Suppl. Fig 2B). Also pointing to a role in 3’UTR regulation, Atx2 edit sites were particularly prevalent in brain transcripts with longer 3’UTRs that are more often subject to translational control (Miura et al., 2013; Wang and Yi, 2014) (Suppl Fig 2C). These observations are consistent with prior work showing that Atx2 can mediate activation or repression of specific mRNA translation via elements in their 3’UTRs (Lee et al., 2017; Lim and Allada, 2013; McCann et al., 2011; Sudhakaran et al., 2014; Zhang et al., 2013).

**Figure 2:**
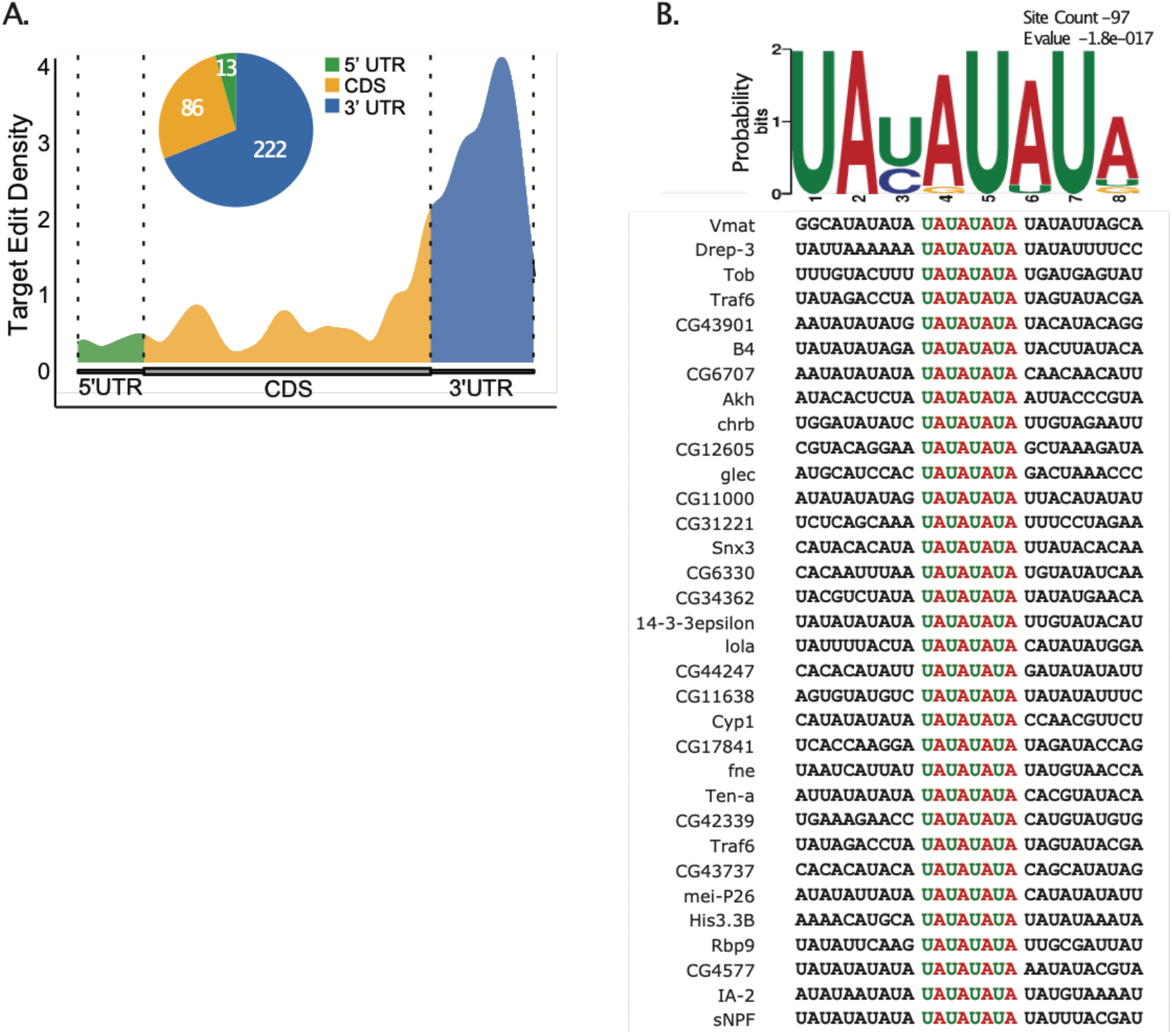
Atx2 preferentially edits AU rich sequences in the 3’UTRs of the target mRNAs. (A) Metaplot analysis showing Atx2 preferentially associates with 3’UTRs of target mRNA. (B) Motif analyses involving +/− 100 bases around the edit site using MEME identifies ARE sequence in the 3’UTRs of target mRNAs.

Further analyses using the MEME suite of tools for motif-based sequence analysis identified an AU-rich element (ARE) ‘UAUAUAUA’ as highly enriched in mRNA target sequences within 100 bases of identified edit sites (Figure 2B). These AREs, previously implicated in regulation of mRNA stability, are most abundant near 3’UTR edits sites of the target mRNAs, while a secondary motif with GC rich sequence was found predominantly for the CDS edits (Suppl. Figure 2D). This suggests a model in which Atx2 preferentially associates with ARE-containing 3’UTRs of target mRNAs, to regulate their stability *in vivo*.

### Atx2 stabilizes the majority of its mRNA targets

AREs are destabilizing motifs on mRNAs that signal their rapid degradation (Vasudevan and Peltz, 2001). For this reason, several RBPs modulate mRNA stability by binding to AREs, and by regulating their accessibility to RNA degradative machinery (Mayya and Duchaine, 2019). For example, Pumilio binding to ARE increases degradation, while HuR binding stabilizes the mRNAs (López De Silanes et al., 2004; Weidmann et al., 2014). To ask how Atx2 binding alters target mRNA stability, we expressed a previously validated Atx2-targeting RNAi construct in fly brains to reduce endogenous Atx2 expression and used RNAseq to determine how this affected steady state levels of Atx2-target and non-target mRNAs (McCann et al., 2011; Sudhakaran et al., 2014).

Experimental *elav-Gal4, UAS-Atx2-RNAi; Tub-Gal80^ts^* flies were reared to adult hood at 18°C. They were then transferred to 29°C to inactivate Gal80^ts^ and enable neural Atx2-RNAi expression for 5 days, before isolating brain RNA (Figure 3A). RNAseq data confirmed Atx2 mRNA knockdown in experimental flies expressing Atx2-RNAi (Figure 3B).

**Figure 3:**
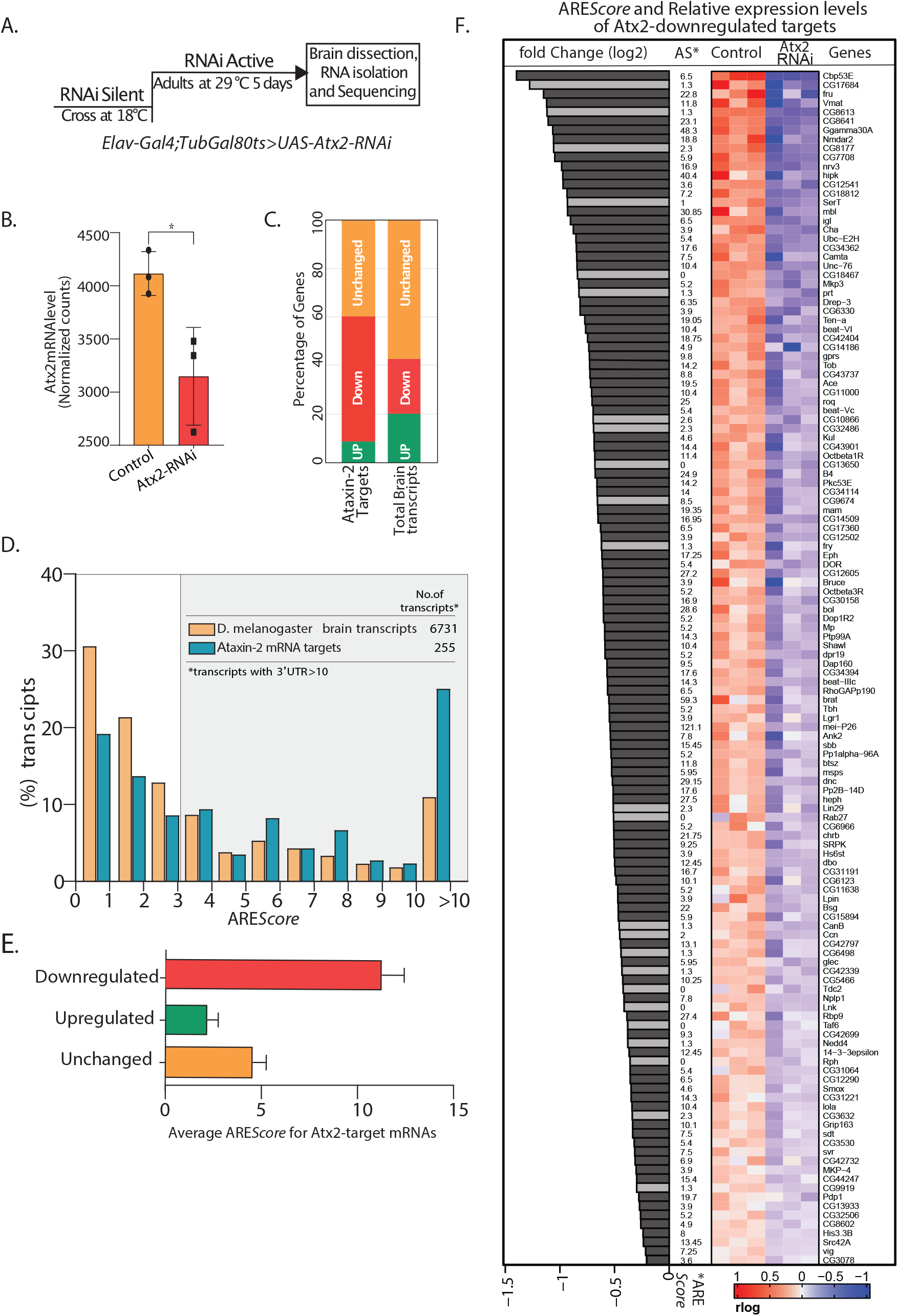
Atx2-ARE interactions modulate mRNA stability. (A) Schematic of strategy to induce RNAi expression specifically in adult fly brain. Total RNA extracted from brain post 5 days at 29°C was sequenced using Illumina 2500 and differential expression analyzed using DESeq2. (B) Normalized mRNA read-counts showing Atx2 mRNA levels to be significantly reduced following RNAi expression compared to Gal4 control with a p-value of <0.0298 using student’s t-test. (C) Effect on Atx2 target mRNAs following Atx2-knockdown: the majority are reduced in level indicating a role for Atx2 is target stabilization. (D) ARE*Score* analysis showing AREs to be enriched in Atx2 target mRNAs compared to the brain transcriptome. (E) Among Atx2-target mRNAs, higher ARE scores are seen in mRNAs whose levels are reduced following Atx2 knockdown. (F) Correlation between ARE*Scores* and relative expression levels for three biological replicates of Atx2-downregulated targets (selected based on with a p-value <0.05). Heatmap indicates relative expression levels for each mRNA in control and experimental brain samples. Light gray bars represent Atx2 target mRNAs without significant ARE*Scores.*

Atx2-knockdown caused a significant reduction in levels of over 53.2% of the Atx2-target mRNAs indicating a broad role for Atx2 in target-mRNA stabilization. This was however not universal, and levels about ~8.8% of the target mRNAs were increased with Atx2 knockdown. Non-target mRNAs were substantially less affected: ~22.5% of all mRNAs from the global brain transcriptome were reduced by Atx2 knockdown, with ~57.2% showing no detectable change in expression levels (Figure 3C). Together, the RNAseq data: (a) provide additional evidence in support of *in vivo* interactions between Atx2 and target-mRNAs identified by TRIBE analysis; and (b) are consistent with Atx2 binding to the ARE motif acting predominantly to stabilize target mRNAs.

ARE*Score* is a numerical assessment of ARE strength with high scores correlating to reduced RNA stability in reporter assays (Spasic et al., 2012). In support of a broad role for Atx2 in regulation of ARE function in neurons, UTRs with high ARE*Scores* are clearly enriched in the Atx2 target mRNAs compared to the general neural transcriptome (Figure 3D). Further, within the group of Atx2 target RNAs identified by TRIBE, higher ARE*Scores* strongly predict mRNAs whose steady-state levels are reduced by RNAi-mediated Atx2 knockdown (Figure 3E-F). These results indicate that Atx2 stabilizes a subset of its target mRNAs by binding to AREs.

The data also point to alternative and/or context-specific mechanisms for mRNA-regulation by Atx2. For instance, some Atx2 target mRNAs lack AREs and a significant subset of ARE containing target mRNAs are not destabilized by Atx2 knockdown. These could be explained by alternative pathways by which Atx2 is recruited to mRNAs, for example via microRNA pathway components (McCann et al., 2011; Sudhakaran et al., 2014), or by further layers of regulation conferred by additional RBPs recruited onto target mRNAs.

To further understand mechanisms by which Atx2 interacts with its target mRNAs, we asked which domains of Atx2 might be required for this function. In particular, we asked whether the unstructured IDR or structured Lsm/LsmAD domains contributed to the specificity of Atx2-mRNA target interactions.

### The cIDR domain enables and the Lsm domain opposes Atx2-interactions with target mRNA

Ataxin-2 has three structured domains (Lsm, LsmAD and PAM2) embedded within extended, poorly structured regions of the protein. Although, interactions mediated by structured domains *in vivo* are necessary for normal organismal viability in *Drosophila*, the mechanism by which structure domains function is only clear for PAM2 which, by binding to polyA binding protein PABP, likely allows interactions with polyA tails of mRNAs (Satterfield and Pallanck, 2006). In contrast, the most prominent disordered regions in Ataxin-2 (mIDR and cIDR) contribute little to animal viability but are selectively required for the assembly of mRNPs in neurons or cultured cells (Bakthavachalu Huelsmeier et al., 2018). To test how Lsm, LsmAD and cIDR domains of Atx2 contribute to the specificity of mRNA-target interactions, we performed TRIBE analyses with specific domain-deleted forms of Atx2. Because purified Lsm domain alone as well as Lsm+LsmAD domains have been reported to bind the AU rich sequences *in vitro*, we generated transgenes expressing a construct with Lsm+LsmAD domains of Atx2 fused to the catalytic domain of ADAR (Yokoshi et al., 2014) (Figure 4A). In addition, we created transgenic lines expressing Atx2 deleted for either Lsm, LsmAD, mIDR or cIDR domains of Atx2, based on the UAS-Atx2-ADARcd construct scaffold. Using the same approach as earlier, we analyzed how each of the domain deletions affected the Atx2-target binding in adult neurons.

**Figure 4:**
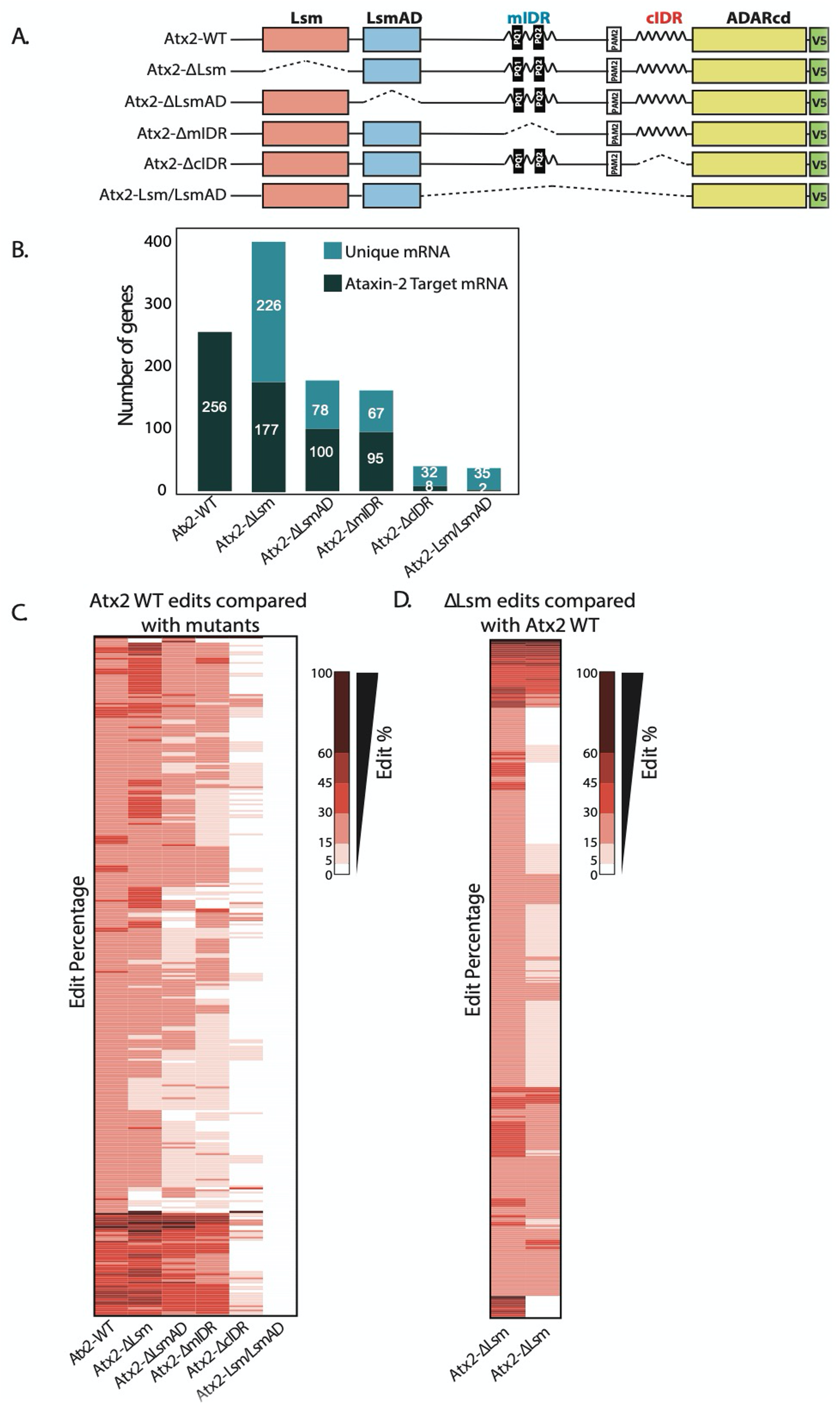
Atx2 requires its cIDR domain for interaction with its target mRNA. (A) An illustration of domains of Atx2. The structured (Lsm and LsmAD) and disordered domains (mIDR and cIDR) of Atx2 protein were deleted one at a time to understand the domains necessary for its interaction with target RNA. The wild type Atx2 and the deletions contained a c-terminal ADARcd and V5 tag. The deletions are shown with the dotted lines. (B) Deletion of LsmAD or mIDR domain reduced the ability of Atx2 to interact with its targets, while cIDR deletion almost entirely prevented Atx2 mRNA interactions. The Lsm-domain deletion showed an overall increase in mRNA edits. (C) More detailed heatmap view of individual target-mRNAs showing target edit percentages in flies expressing control or domain-deleted forms Atx2 fused to ADARcd. (D) Most of the apparently new targets identified in Atx2ΔLsm TRIBE analyses are also edited, at with lower efficiency, in flies expressing full-length Atx2-ADAR fusions. This suggests that deletion of the Lsm domain increases interaction of Atx2 with its native target mRNAs. (For C and D, 15% threshold edits identified for samples in column 1 (Atx2WT for C and Atx2ΔLsm in D) was used to compare with all the edits of rest of the columns to generate a heatmap).

Initial observations indicated that Lsm+LsmAD domains on their own could not efficiently target ADAR to mRNA: transcripts sequenced from Lsm+LsmAD-ADARcd expressing brains contained negligible edits, which did not overlap significantly with Atx2 targets. Therefore, the Lsm and LsmAD domains are insufficient to drive Atx2-mRNA interactions *in vivo*. Further, deletions of either Lsm or LsmAD did not block the ability of Atx2 to interact with most TRIBE targets: thus, they appear neither necessary nor sufficient for Atx2 targeting to these mRNAs (Figure 4B and C). These surprising observations led us to examine the role of disordered domains in driving Atx2-mRNA interactions *in vivo*.

In contrast to effects of deleting the Lsm domain, deletion of cIDR, abolished Atx2 binding to most target mRNAs (Figure 4B and C). As the cIDR plays a major role in mRNP assembly, this unexpected observation suggests Atx2 moves into proximity of target mRNAs only after cIDR-mediated granule formation. Deletion of the mIDR resulted only in a relatively minor reduction in target binding, but this is consistent with prior work indicating only a minor role in mRNP assembly (Bakthavachalu, Huelsmeier et al., 2018). Deeper analyses provided additional support for a model in which Atx2 associates to RNA-binding proteins in individual mRNPs, but is brought into closer contact with mRNAs through remodeling events associated with the formation of higher order mRNP assemblies.

A key observation is that while Lsm-domain deletions did not reduce RNA edits, they also curiously resulted in a significantly larger number of edited target mRNAs compared to the full length, wild-type Atx2 (Figure 4C). A more detailed analysis showed that several of the apparently novel targets of Atx2ΔLsm were also bound by the wild type Atx2 but with reduced affinity and were therefore below the threshold of our analysis (Figure 4D and Suppl. Figure 3C). Because sequencing depth, read quality and ADAR mRNA levels were similar across control and domain-deletion experiments (Suppl. Figure 3A), these observations argue that the Lsm domain acts to broadly antagonize physiologically relevant mRNA interactions driven by the cIDR. Thus, while the cIDR domain of Atx2 is essential for its mRNA target interactions, the Lsm domain is inhibitory: in its absence, Atx2 shows enhanced association with its native, target mRNAs.

### The Atx2 Lsm domain inhibits cIDR mediated mRNP granule assembly

The most parsimonious explanation for the observed opposing effects of Lsm and cIDR domain deletions on mRNA editing (Figure 4), is that the Lsm domain functions to oppose cIDR mediated mRNP-granule assembly (Bakthavachalu, Huelsmeier et al., 2018). To directly test this hypothesis, we first asked whether deletion of the Lsm domain enhanced Atx2 RNP assembly. Expression of wild type C-terminally SNAP tagged Atx2 (Atx2-SNAP) in S2 cells led to the formation of SG like RNP-granule foci through a mechanism dependent on cIDR as previously described (Figure 5(ii and v)) (Bakthavachalu, Huelsmeier et al., 2018). Strikingly consistent with our predictions, expression of Atx2ΔLsm-SNAP constructs lacking the Lsm domain induced significantly larger RNP granules as compared to wild type Atx2-SNAP (Figure 5(i)). Similar to wild type Atx2-SNAP granules, these large Atx2ΔLsm-SNAP induced granules also contained the SG protein, Rasputin (Rin) (Suppl. Figure 4) and their formation required the presence the cIDR (Figure 5(iv)). Thus, Lsm domain appears to act antagonistically to cIDR to prevent RNP granule assembly.

**Figure 5:**
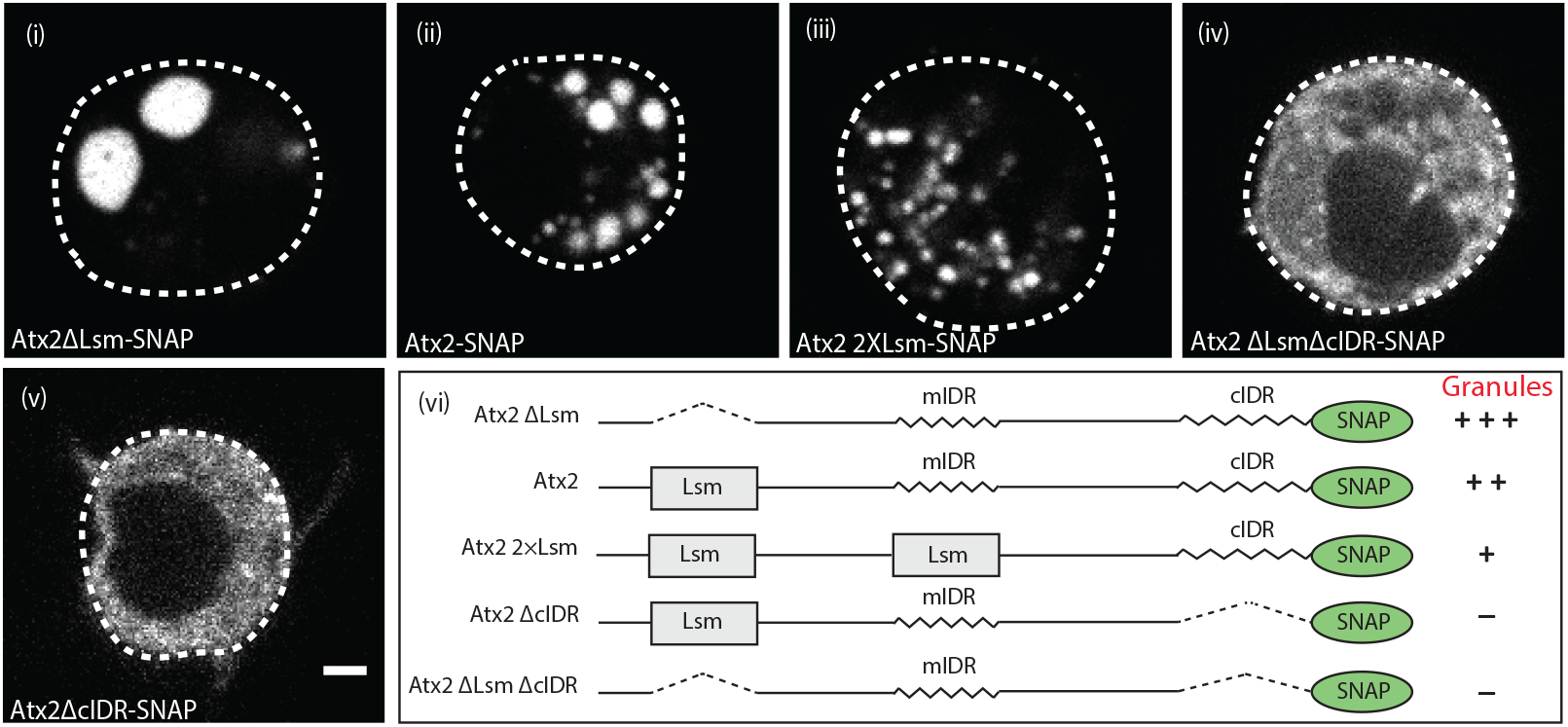
Atx2 Lsm domain alters granule dynamics. *Drosophila* S2 cells expressing Atx2 protein with a C-terminal SNAP tag. Atx2ΔLsm form large cytoplasmic granules (i) which is much larger than WT Atx2 granules (ii). Atx2 with an additional Lsm domain in place of mIDR to create 2XLsm forms smaller granules compared to WT Atx2 in S2 cells (iii). Deletion of cIDR domain blocks Atx2 granule formation (iv). In the absence of cIDR, Lsm deletion does not rescues Atx2 granules (v). Domain map along with the granule phenotype is shown in (vi). The scale bar corresponds to 2µm.

To further confirm this, we tested if the inclusion of additional Lsm domains would reduce the ability of Atx2 to form RNP granules. As the deletion of the Atx2 mIDR does not alter mRNP granule assembly in S2 cells (Bakthavachalu, Huelsmeier et al., 2018), we replaced mIDR with an additional Lsm domain to create an Atx2 protein with two Lsm domains. Remarkably, Atx2 with two Lsm showed much smaller foci in S2 cells (Figure 5(iii)). The summary of these results is shown in Figure 5(vi). Taken together, the observations that deletion of the Lsm enhances and addition of an extra Lsm domains inhibits RNP assembly provide strong support for two opposing activities encoded by the Lsm and cIDR domains of Atx2.

## DISCUSSION

Previous work has shown that *Drosophila* Atx2 functions in neurons as a translational activator of the *period* mRNA that controls circadian rhythms, as a translational repressor of the calcium-calmodulin dependent kinase CaMKII involved in synaptic plasticity and memory (Lim and Allada, 2013; Sudhakaran et al., 2014; Zhang et al., 2013). Atx2 is also required for assembly of neuronal mRNPs believed to provide pools of synaptically localized mRNAs whose translation contributes the consolidation of long-term memory (McCann et al., 2011). These studies indicate the specific positive and negative translational functions of Atx2, which are mediated by structured-domain interactions with Lsm12 or Me31B/DDX6, respectively (Lee et al., 2017), and that mRNP-assembly functions are mediated by its cIDR (Bakthavachalu, Huelsmeier et al., 2018). However, the generality of these mechanisms, the range of neuronal mRNAs and neuronal functions under Atx2 regulation, as well as how structured and disordered domain interactions are coordinated, remains largely unknown. Here, by deploying and building on TRIBE analysis to identify a suite of Atx2-target mRNAs, we address these questions and provide insights of relevance for biology, technology and medicine.

### Neural functions of Ataxin-2

TRIBE allows *in vivo* RNA targets of RBPs to be identified from small tissue samples, eliminating several technical challenges and artifacts associated with immunoprecipitation based methods (Jin et al., 2020; McMahon et al., 2016; Xu et al., 2018). This method led to identification of 256 *Drosophila* brain mRNAs that associate with Atx2 with the proximity and stability required for Atx2-linked enzymatic editing of the mRNA. These mRNAs are reproducibly identified in replicate experiments and do not show any over-representation of highly expressed mRNAs. Moreover, the observation that a substantial fraction of these mRNAs either have AREs in their 3’UTRs and/or show altered steady-state levels following Atx2 knockdown, argues that the majority represent real Atx2-targets and not non-specific proximity-based editing events that can sometimes occur within RNP complexes (Biswas et al., 2019). Thus, the resulting robust dataset of Atx2 targets may provide valuable hypotheses for biological functions and genetic pathways regulated by Atx2. For instance, a striking enhancement of mRNAs encoding specific neuropeptides and neuronal hormones suggests that altered intercellular communication mediated by their translational regulation may contribute to behavioral plasticity associated with circadian time or long-term memory. Similarly, a large subset of target mRNAs encoding proteins regulating neural excitability through multiple different mechanisms is unexpected, and points to the possibility that activity-regulated translation may act via local changes in membrane properties to achieve localized plasticity required for encoding specific memories.

But not all Ataxin-2 regulated mRNAs have been identified. It is notable that two of the best-established Atx2-targets, CaMKII and *per* were not identified by TRIBE. While the *per-*Atx2 interactions, being time- and cell-type restricted (Lim and Allada, 2013; Zhang et al., 2013), could potentially be missed for statistical reasons, this is unlikely the case for CaMKII, a highly expressed mRNA that co-immunoprecipitates with Atx2 (Sudhakaran et al., 2014). We suggest instead, that these represent Atx2 targets missed by TRIBE, because they are regulated through relatively indirect mechanisms that do not require close contact between Atx2 and the mRNA. For instance, in case of *per*, its 3’UTR is recognized by the sequence-specific RBP Twentyfour (TYF), which recruits Atx2 that in turns recruits a Lsm12-containing complex to the *per* 3’UTR thus allowing translational activation (Lee et al., 2017). Similarly, for CaMKII, Atx2 may be recruited by miRNA pathway components and act via co-regulators such as Me31B/DDX6, through mechanisms that do not require direct contact between Atx2 and target mRNAs. The above may also help explain why previously proposed target mRNAs in metabolic pathways for instance are not represented in this dataset (Yokoshi et al., 2014). Indeed, Atx2 likely binds to several additional neuronal mRNAs not identified by TRIBE, which requires Atx2-proximity to the mRNA. Such targets may be better identified by CLIP-based methods. However, considerable new understanding can be provided by the detailed analysis of the 256 robust targets identified here by TRIBE.

One important insight is the discovery of a broad function for Atx2 in neuronal mRNA stabilization. Atx2 associates preferentially to 3’UTRs of the target-mRNAs, and particularly to AU-rich sequences (AREs) in these UTRs (Figure 2). AREs are common cis-regulatory features of rapidly degrading mRNAs, a posttranscriptional gene regulation strategy adopted by all eukaryotic cells (García-Mauriño et al., 2017). The observation that RNAi-based knockdown of Atx2 in *Drosophila* brain causes levels of a large percentage of the Atx2 target mRNAs to be significantly reduced (Figure 3B, C) and that the most downregulated targets have strong AREs (Spasic et al., 2012) suggests that Atx2 directly or indirectly associates with AREs to protect mRNAs from degradation (Figure 3E, F). This could be achieved by blocking ARE-dependent recruitment of RNA degradation complexes, through a mechanism similar to that described previously for HuR (Peng et al., 1998). These conclusions may also be relevant for mammalian Atxn2, as physical interactions between mammalian Atxn2 and AREs have been previously described using PAR-CLIP analyses from cultured HEK293 cells (Yokoshi et al., 2014). Moreover, Atxn2-CAG100-KnockIn mouse engineered to express polyQ expanded forms of Ataxin-2 that should enhance granule formation, show a predominant upregulation of mRNAs, consistent with a role for Atx2-mediated mRNP assembly in stabilizing target mRNAs (Sen et al., 2019).

It is important to note that some mRNAs with high ARE scores do not appear to be stabilized by Atx2 and conversely some that do not contain AREs appear to be affected by Atx2 knockdown (Fig. 3E). Both of these observations are consistent with additional layers and mechanisms of regulation conferred by co-regulating RBPs: either by providing alternative pathway for ARE regulation via for instance miRNA binding (Sun et al., 2009; Van Kouwenhove et al., 2011), or an alternative mechanism for recruitment of Atx2 to the 3’UTR of mRNAs.

### Mechanisms of Ataxin-2 function in RNP granule assembly

Our work provides two insights into the mechanisms of mRNP formation. First, it indicates that individual mRNPs may be substantially remodeled as they assemble into higher order mRNP assemblies. In support of this, we show that Atx2 lacking its cIDR, which cannot form granules, is also not associated closely enough with mRNAs to allow their editing by a linked ADAR catalytic domain. One possibility is that mRNP remodeling is driven by major conformational changes in RBPs, which not only increase their propensity to drive mRNP condensation, but also result in altered RBP-RBP and RBP-RNA interactions. In this context, recent work on G3BP/Rin has shown that the protein exists in two dramatically different conformational states: a closed form, in which its IDRs are inaccessible for condensation reactions, and a dephosphorylation-induced open form, capable of mediating SG association (Guillén-Boixet et al., 2020; Laver et al., 2020; Sanders et al., 2020). In such a framework, it is easy to see how Atx2 interactions with RBPs and mRNAs could be altered under conditions that support granule assembly. While these changes in Atx2 interactions could occur due to structural changes in other components of Atx2-containing mRNPs, our second insight is that alterations in the Atx2 protein itself probably occur and contribute to driving granule assembly.

Ataxin-2 is a modular protein capable of association with multiple translational control components (Dansithong et al., 2015; Lastres-Becker et al., 2016; Lee et al., 2017; Satterfield and Pallanck, 2006; Swisher and Parker, 2010). Although Atx2 lacks RNA recognition domains like RRMs, KH or other previously characterized RNA-binding domains, homology-based modeling studies and indirect experimental observations have suggested that the Lsm domain of Atx2 may mediate RNA interaction (Calabretta and Richard, 2015; Hentze et al., 2018; Yokoshi et al., 2014). However, direct experimental tests of this hypothesis show that close Atx2 interactions with mRNA, as assessed by TRIBE, are actively prevented by the Lsm domain, which also opposes mRNP assembly (Figure 4 and 5). In contrast, the cIDR which drives mRNP assembly is necessary for Atx2-coupled editing of target mRNA.

The simplest explanation for these findings is that Atx2-association with individual, potentially translationally active mRNPs in the soluble phases are mediated by Lsm domain-RBP interactions that also occlude or prevent cIDR-mediated mRNP assembly (Ciosk et al., 2004; Lee et al., 2017; Satterfield and Pallanck, 2006). Conditions that promote mRNP assembly disrupt Lsm-domain mediated interactions and enable cIDR-driven granule formation (Figure 6). We note that recent work on G3BP has beautifully elaborated phosphorylation-regulated intramolecular interactions that similarly allow the molecule to switch between soluble and assembly competent conformations (Guillén-Boixet et al., 2020; Laver et al., 2020; Sanders et al., 2020). Though our experiments do not yet define molecular and biophysical details by which Atx2 transitions from assembly-inhibited to assembly-competent states, our observations: (a) clearly demonstrate crucial opposing, physiological roles of the Lsm and cIDR domains in this process; and (b) suggest that regulation of intermolecular interactions mediated by the Lsm domain will be involved in control of Atx2-mediated granule assembly.

**Figure 6:**
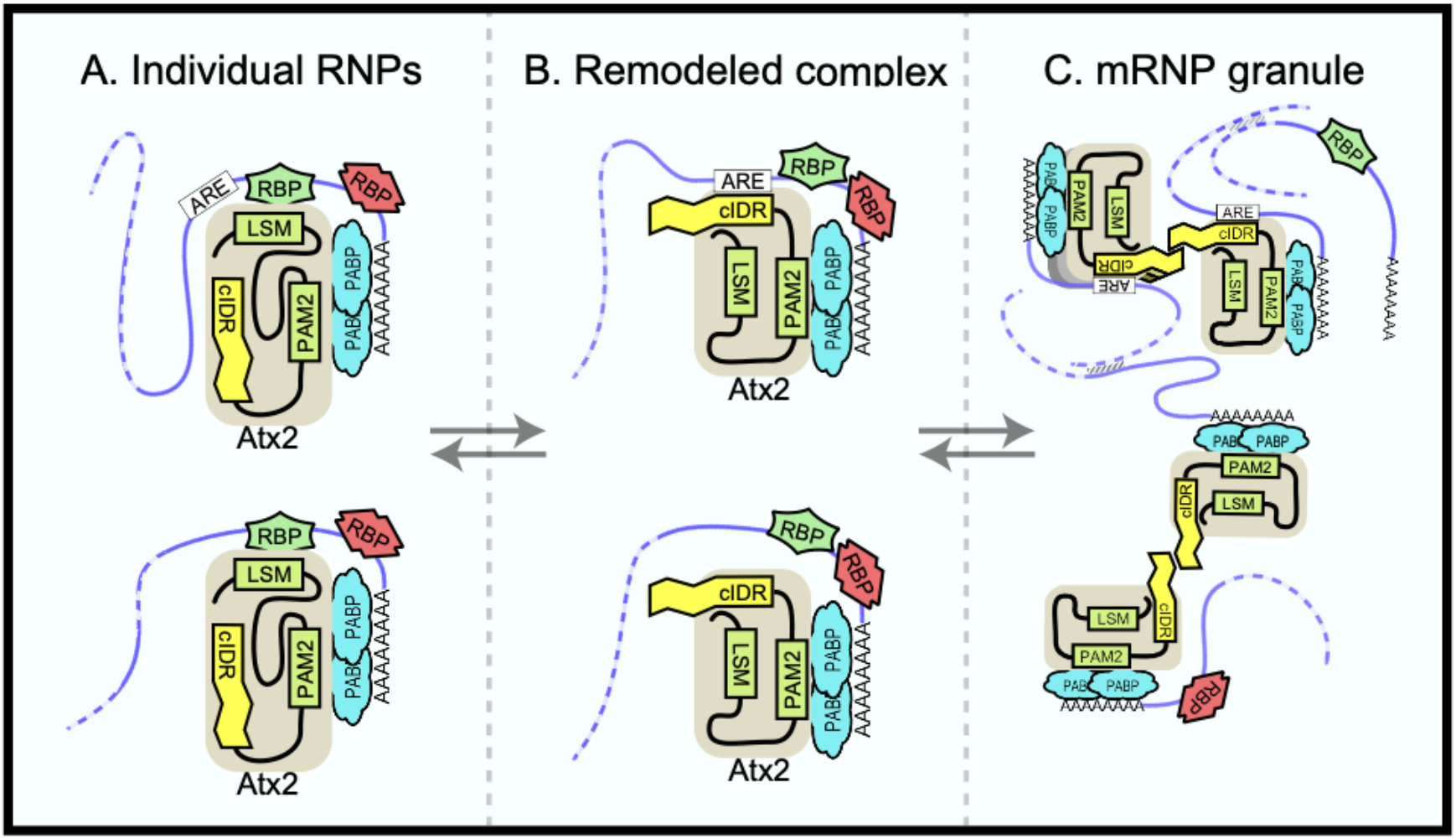
Ataxin-2 domains in mRNA target regulation. A model to explain current and previous observations on Ataxin-2 as well as its function in mRNA regulation. (A) Ataxin-2 does not make direct contact with mRNAs in soluble mRNPs. Instead, it is recruited to RNA by other RBPs that bind to structured Lsm (or LsmAD) domains. In these soluble mRNPs, the cIDR is buried and inaccessible. One class of mRNP (above) contain AREs; the second class below (e.g. *per* in *Drosophila*) does not. (B) Under specific signaling conditions (e.g. stress) RBP-Atx2 interactions are prevented. In these “remodeled” mRNPs, the Atx2-cIDR is exposed. We speculate that that a section of this IDR directly binds to nearby AREs, while other sections mediate multivalent interactions that contribute to mRNP condensation. (C) Ataxin-2 cIDR interactions enable mRNP assembly into granules, facilitated by RNA-RNA crosslinks and interactions mediated by other RBPs (e,g. G3BP/Rin). These RNP granules may include both ARE containing mRNAs (edited) as well as mRNA those that don’t. Note that additional RBPs (red) could determine not only Atx2 proximity to specific mRNAs (leading to editing), but also the effect of Atx2 on mRNA translation and/or stability. These and alternative models remain to be tested.

It is important to note that Ataxin-2 has additional, essential functions beyond those described here. In particular, given Atx2 structured domains not required for TRIBE-target binding are essential for survival, unlike the IDR, which is required for editing of TRIBE targets but not for animal survival, it appears likely that a class of Atx-2 target mRNAs is regulated outside of mRNP granules though largely structured-domain interactions (Figure 6). Additional approaches and experiments are required to identify such mRNAs as well as mechanisms by which they are regulated.

### Insight for disease and therapeutics

Ataxin-2 has attracted considerable clinical interest for three main reasons. First, assembly promoting mutations in the Ataxin-2 gene or associated RNA-binding and SG proteins such as TDP-43 can cause neurodegenerative disease. Second, SG proteins such TDP-43 are usually present in cytoplasmic protein inclusions associated with familial and heritable forms of ALS and FTD. And third, reduction of Ataxin-2 can slow neurodegeneration in animal models of ALS, indicating that the normal function of Ataxin-2 is required for initiation or progression of disease (Becker et al., 2017; Scoles et al., 2017). These findings, interpreted in a framework wherein SG are thought facilitate the nucleation of pathogenic amyloid filaments, have led to development of therapeutics based on reducing levels of Ataxin-2, for example using antisense oligonucleotides (ASOs) by major companies such as Ionis and Biogen Inc. In this context, the discovery that the Lsm domain inhibits mRNP assembly suggests, first, that mutations inactivating this domain could have effects similar to polyQ expansions and promote disease and second, that compounds targeting specific domains and activities of Ataxin-2 may prove more effective as therapeutics than those which knock down protein levels.

The case for our understanding of the function of each Atx2 domain and developing-specific modulators is particularly strong since Ataxin-2 protein itself has several important roles not only in mRNA stabilization as shown here, but also in protein translation, cell signaling, metabolism and embryonic development (Halbach et al., 2017; Kato et al., 2019; Lim and Allada, 2013; Meierhofer et al., 2016; Yang et al., 2019; Yokoshi et al., 2014; Zhang et al., 2013) as shown by various biological studies of native Ataxin-2 function.

## ACKNOWLEDGEMENTS

We thank Roy Parker and members of the Ramaswami, VijayRaghavan and Bakthavachalu labs for useful discussions and/or comments on the manuscript. Thanks to Michael Rosbash for *Drosophila* TRIBE plasmid and to colleagues acknowledged in the Key Resources table for generously sharing essential reagents and informal advice. The fly facility at Bangalore Life Science Cluster (BLiSC) provided support with fly stock supply as well as generation of transgenic; CIFF at BLiSC provided essential confocal microscopy support; and Awadhesh Pandit and next generation genomics facility at BLiSC provided NGS service. We acknowledge Drosophila Genomics Resource Centre (supported by NIH grant 2P40OC010949) for *Drosophila* S2 cells.

## AUTHOR CONTRIBUTIONS

Conceptualization, A.S., J.Huelsmeier, K.V.R., M.R., and B.B.; Methodology, A.S., J.Huelsmeier, A.R.K., S.S.P., J.Hillebrand, A.P., D.T., R.T.K., K.A., K.V.R., M.R., and B.B.; Investigation, A.S., J.Huelsmeier, A.R.K., S.S.P., J.Hillebrand, A.P., D.T., R.T.K., K.A., K.V.R., M.R., and B.B.; Writing–Original Draft, A.S., J.Huelsmeier, K.V.R., M.R., and B.B.; Writing–Review & Editing, A.S., J.Huelsmeier, A.R.K., S.S.P., J.Hillebrand, A.P., D.T., R.T.K., K.A., K.V.R., M.R., and B.B.; Funding Acquisition, A.S., K.V.R., M.R., and B.B.; Resources, Fly community.

## EXPERIMENTAL MODEL AND SUBJECT DETAILS

### Generation and rearing of Drosophila stocks

Drosophila stocks were maintained at 25°C in corn meal agar and experimental fly crosses were done as specified in the respective experimental methods. List of Drosophila stocks used, and transgenic flies generated for this study are listed in Suppl. Table 2.

### S2 cell culture

Drosophila S2R+ cells were obtained from DGRC and cultured in Schneider’s medium with 10% FBS, penicillin and streptomycin at 25°C.

## METHODS DETAILS

### Creation of transgenic animals

*Drosophila* Atx2 full length cDNA was cloned into pJFRC7-20XUAS-IVS-8_ADARcd plasmid (a gift from Rosbash lab) to create pJFRC7-20XUAS-IVS-8_Atx2wt-ADARcd plasmid. Domain deletions were created using overlapping PCR and Gibson assembly or non-overlapping PCR and ligation using pJFRC7-20XUAS-IVS-8_Atx2wt-ADARcd as template. Sequence confirmed plasmids were used to generate transgenic *Drosophila* using PhiC31integrase dependent site-specific insertion of the transgene on the second chromosome. Details of plasmids used for transgenesis are listed in Suppl. Table 3. Embryo injections were performed at NCBS transgenic fly facility. Primers used for domain deletions are listed in Suppl. Table 4.

### Experimental fly crosses

Strains homozygous for the elav-GAL4 and tub-Gal80^ts^ transgenes were crossed with homozygous UAS-transgenic flies at 18°C till the adult fly emerged. One day old adult flies from the crosses were maintained at 29°C for 5 days before processing for RNA extraction.

### RNA extraction from brain and NGS

Total RNA was isolated from adult brain (10-12 brains per replicate) dissected in RNA Later using TRIzol reagent (Invitrogen) as per the manufacturer’s protocol. Illumina libraries were prepared from Poly(A) enriched mRNA using NEBNext® Ultra™ II Directional RNA Library Prep kit (E7765L) or TruSeq RNA Library Preparation Kit V2 (RS-122-2001) and sequenced with Illumina HiSeq 2500 system. Atx2-wt TRIBE samples were sequenced using HiSeq SBS Kit v4 (FC-401-4003) producing 2X125 paired end non-strand specific reads. TRIBE for all the Atx2 domain mutants were sequenced using HiSeq PE Rapid Cluster Kit v2 (PE-402-4002) to generate 2X100 paired end strand-specific data.

### TRIBE Data Analysis

All sequencing reads obtained post adaptor removal had a mean quality score (Q-Score) >= 37 and so no trimming was required. Sequence details are listed for each experiment in Suppl. Table 5. The TRIBE data analysis was performed as previously described (Rahman et al. 2018) with few modifications. Tools used for analysis are listed in Suppl. Table 6 Briefly, sequencing reads obtained were mapped to dm6 *Drosophila melanogaster* genome using TopHat2 (Trapnell et al., 2009) with the parameters “--library-type fr-firststrand -m 1 -N 3 --read-edit-dist 3 -p 5 -g 2 -I 50000 --microexon-search --no-coverage-search -G dm6_genes.gtf”. Non-strand specific sequencing reads were aligned using tophat2 with the parameters “-m 1 -N 3 --read-edit-dist 3 -p 5 -g 2 -I 50000 --microexon-search --no-coverage-search -G dm6_genes.gtf”. The uniquely mapped SAM output file was loaded in the form of MySQL table with genomic coordinates. Edits for the brain samples were identified by comparing the nucleotide at each position of the genomic coordinates between experiment and control samples and output was printed as bedgraph file. A threshold file was created by ensuring only edits with coverage of at least 20 reads and 15% edits were retained. This threshold file was used for all further analysis unless specified in figure legends.

### Differential Expression data analysis

RNA sequencing reads were mapped to dm6 *Drosophila melanogaster* genome using STAR v2.5.3a with default parameters and read counts were obtained using HTseq with “-s reverse” parameter. DeSeq2 was used for differential expression analysis as previously described (Love et al., 2014).

### ARE analysis

ARE*Scores* tool (http://arescore.dkfz.de/arescore.pl) was used to perform ARE analysis. Only transcripts with 3’UTR > 10 nt in length were considered for analysis. mRNA with highest ARE*Score* was used when multiple transcript variants mapped to the same gene (isoforms of a gene).

### Motif Analysis

The edit coordinates from the bed file were extended by 100bp on either side using Bedtools slop. Intron less sequences with in this +/−100 base pairs were extracted using twoBitToFa. MEME-suite was used to perform motif analysis on the generated FASTA sequences.

### Immunohistochemistry of adult Drosophila brains and S2 cells

Six-day old adult fly brain was dissected in PBS and fixed in PBS containing 4% PFA for 15 mins at room temperature. The brains were than processed for immunostaining according to Sudhakaran et al. (2014). Atx2 ADARcd was stained was using rabbit anti-V5 antibody at 1:200 over one night at 4°C along with neuropil staining using mouse anti-Nc82 (1:100) (Kittel et al., 2006). Secondary antibodies (1:1,000) staining was using anti-rabbit Alexa 488 and anti-mouse Alexa 555 (Molecular Probes) at room temperature for 2 hours. Stained brains were mounted in Vectashield Mounting Medium (Vector Laboratories) and imaged on a Zeiss LSM880 confocal microscope. S2 cells were prepared as described earlier (Bakthavachalu, Huelsmeier et al 2018). In brief, cells were fixed with 4% paraformaldehyde for 10 min at room temperature, followed by permeabilization with 0.05% Triton-X-100 for 10 min. This was followed by blocking with 1% BSA for 30 min. The cells were then stained with primary antibodies against Atx2 (1:500), V5 (1:500) or Rasputin (1:500) followed by probing with corresponding secondary antibodies conjugated with fluorophores. Confocal imaging was done using 60x/1.42 oil objective of Olympus FV3000 microscope. When proteins were SNAP-tagged, SNAP-Surface Alexa Fluor 546 was added after permeabilizing the cells.

### Quantification of Atx2 granules in S2 cells

S2 cells were transfected, fixed and processed as described earlier (Bakthavachalu, Huelsmeier et al 2018). Confocal images were processed using CellProfiler tool (https://cellprofiler.org/) using the watershed algorithm to obtain the size of each granule. At least 80 cells were included in each condition.

### Quantification and statistical analysis

The sample sizes are specified in the figures and figure legends of each experiments. The errors are represented as ± SEM with p-values (*P < 0.05, ****P < 0.0001) calculated by two-tailed Student’s t test and Mann Whitney test. Statistical analysis was performed in GraphPad Prism.

## CONTACT FOR REAGENT AND RESOURCE SHARING

Further information and requests for resources and reagents should be directed to and will be fulfilled by the Lead Contacts, Mani Ramaswami and Baskar Bakthavachalu

## SUPPLEMENTARY MATERIAL

## SUPPLEMENTARY TABLES

**Supplementary table 1:** Reagents used in the study.

| REAGENT or RESOURCE | SOURCE | IDENTIFIER |
| --- | --- | --- |
| **Antibodies** |  |  |
| Chicken anti-Atx2 | Bakthavachalu, Huelsmeier et al 2018 | N/A |
| Rabbit anti-Rasputin | Aguilera-Gomez et al., 2017 | N/A |
| Rabbit anti-GFP | Molecular probes | Cat# A11122 |
| Chicken anti-GFP | Abcam | Cat# mAb 13970 |
| Rabbit anti-V5 | Santa Cruz Biotechnology | N/A |
| Mouse anti-Nc82 | Kittel et al 2006 | N/A |
| Alexa Fluor® 555 goat anti-chicken IgG | Invitrogen | Cat# A21437 |
| Alexa Fluor® 488 goat anti-chicken IgG | Invitrogen | Cat# A11039 |
| Alexa Fluor® 647 goat anti-chicken IgG | Invitrogen | Cat# A21449 |
| Alexa Fluor® 555 goat anti-rabbit IgG | Invitrogen | Cat# A21428 |
| Alexa Fluor® 488 goat anti-rabbit IgG | Invitrogen | Cat# A11078 |
| Alexa Fluor® 647 goat anti-rabbit IgG | Invitrogen | Cat# A21244 |
| Alexa Fluor® 555 goat anti-mouse IgG | Invitrogen | Cat# A21422 |
| Alexa Fluor® 488 goat anti-mouse IgG | Invitrogen | Cat# A21121 |
| Alexa Fluor® 647 goat anti-mouse IgG | Invitrogen | Cat# A21235 |
| **Chemicals, Peptides, and Recombinant Proteins** |  |  |
| Vectashield Mounting Medium | Vector Laboratories | Cat# H-1000 |
| SNAP-Surface 549 | New England Biolabs | Cat# S9112S |

**Supplementary table 2:**
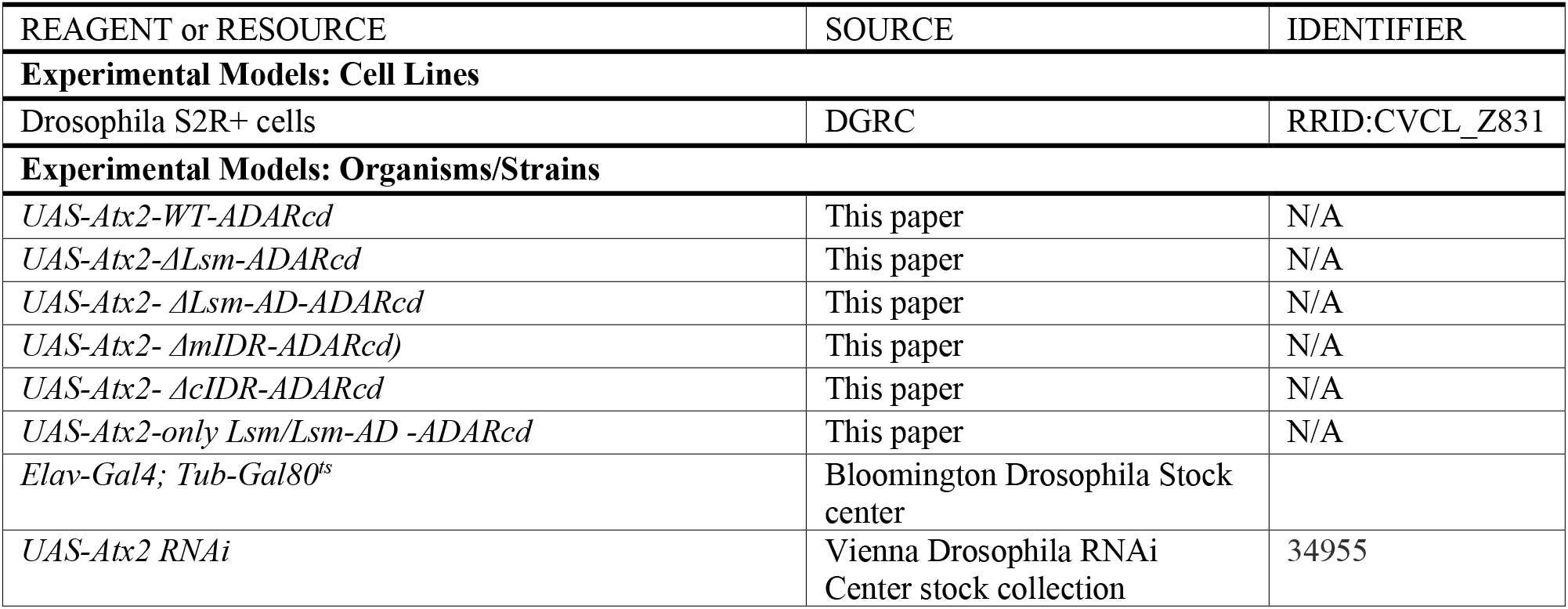
Fly stocks and Cell lines used in the study.

**Supplementary table 3:** Plasmids used in the study

| Recombinant DNA | SOURCE | IDENTIFIER |
| --- | --- | --- |
| pJFRC7-20XUAS-IVS-8 _Atx2-ADARcd | This paper | N/A |
| pJFRC7-20XUAS-IVS-8 _Atx2ΔmIDR -ADARcd | This paper | N/A |
| pJFRC7-20XUAS-IVS-8 _Atx2ΔcIDR -ADARcd | This paper | N/A |
| pJFRC7-20XUAS-IVS-8 _Atx2ΔLsm -ADARcd | This paper | N/A |
| pJFRC7-20XUAS-IVS-8 _Atx2ΔLsmAD -ADARcd | This paper | N/A |
| pJFRC7-20XUAS-IVS-8 _Atx2only-Lsm-LsmAD -ADARcd | This paper | N/A |

**Supplementary table 4:**
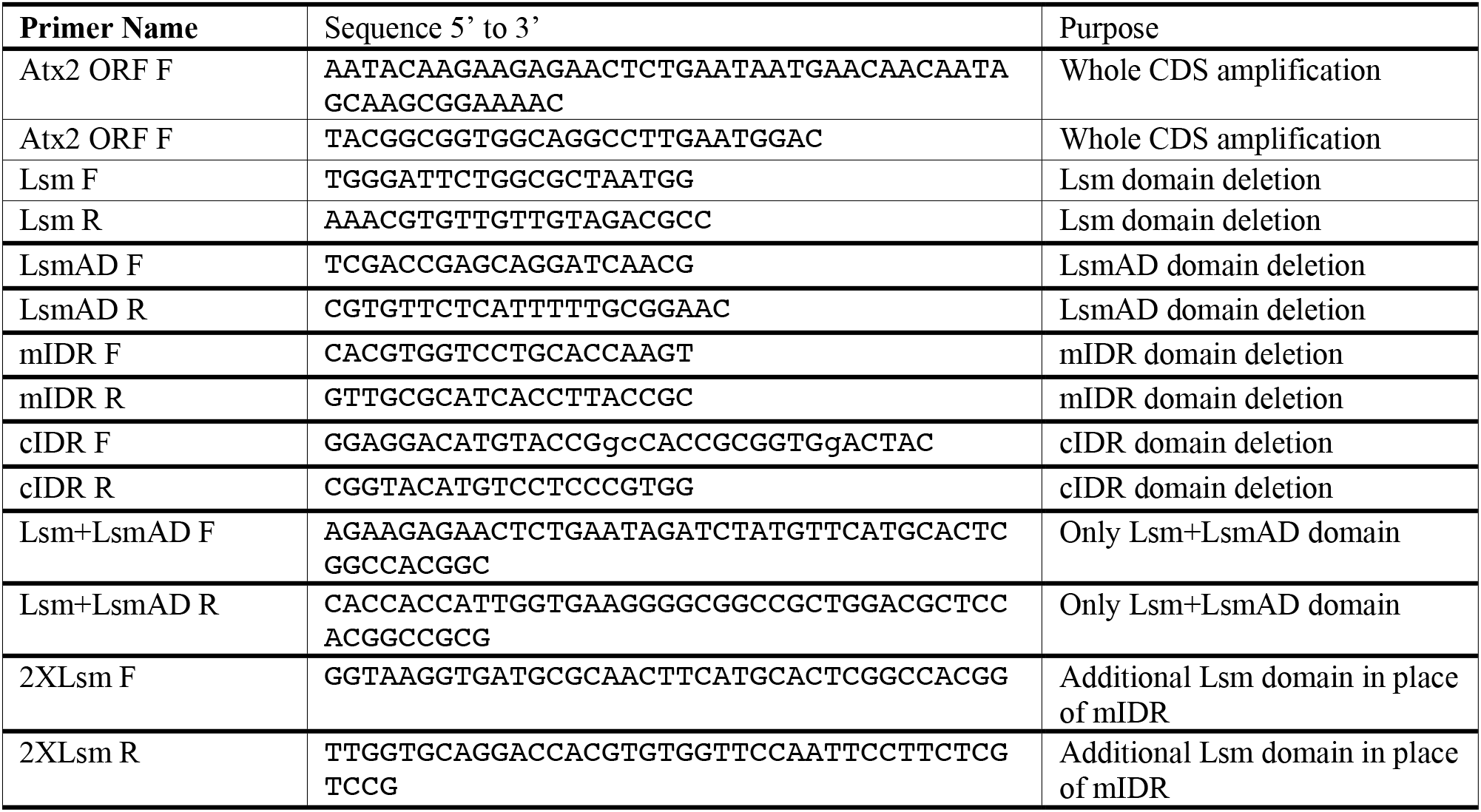
Primers used in the study.

**Supplementary table 5**: The sequencing data and source data are shared as external link. The link for the data repositories will be updated later.

**Supplementary table 6:** Analysis tools used in the study.

| Software and Algorithms | Reference | Source/Link |
| --- | --- | --- |
| TRIBE | McMahon et al., 2016 | https://github.com/rosbashlab/TRIBE |
| STAR v2.5.3 | Dobin et al., 2013 | https://github.com/alexdobin/STAR |
| HTSeq v0.11.2 | Anders et al., 2015 | https://github.com/htseq/htseq |
| DESeq2 | Love et al., 2014 | https://bioconductor.org/packages/release/bioc/html/DESeq2.html |
| AREscore | Spasic et al., 2012 | http://arescore.dkfz.de/arescore.pl |
| Guitar | Cui et al., 2016 | https://bioconductor.org/packages/release/bioc/html/Guitar.html |
| Bedtools | Quinlan and Hall, 2010 | https://github.com/arq5x/bedtools2 |
| twoBitToFa | - | https://genome.ucsc.edu/goldenPath/help/twoBit.html |
| MEME suite | Bailey et al., 2009 | http://meme-suite.org/tools/meme |
| Cellprofiler | McQuin et al., 2018 | https://cellprofiler.org |
| ImageJ | Schneider et al., 2012 | https://imagej.nih.gov/ij/ |
| Ggplot2 | Wilkinson, 2011 | https://github.com/tidyverse/ggplot2 |
| Pheatmap |  | https://cran.r-project.org/web/packages/pheatmap/index.html |

## SUPPLEMENTARY FIGURES

**Supplementary Figure 1:**
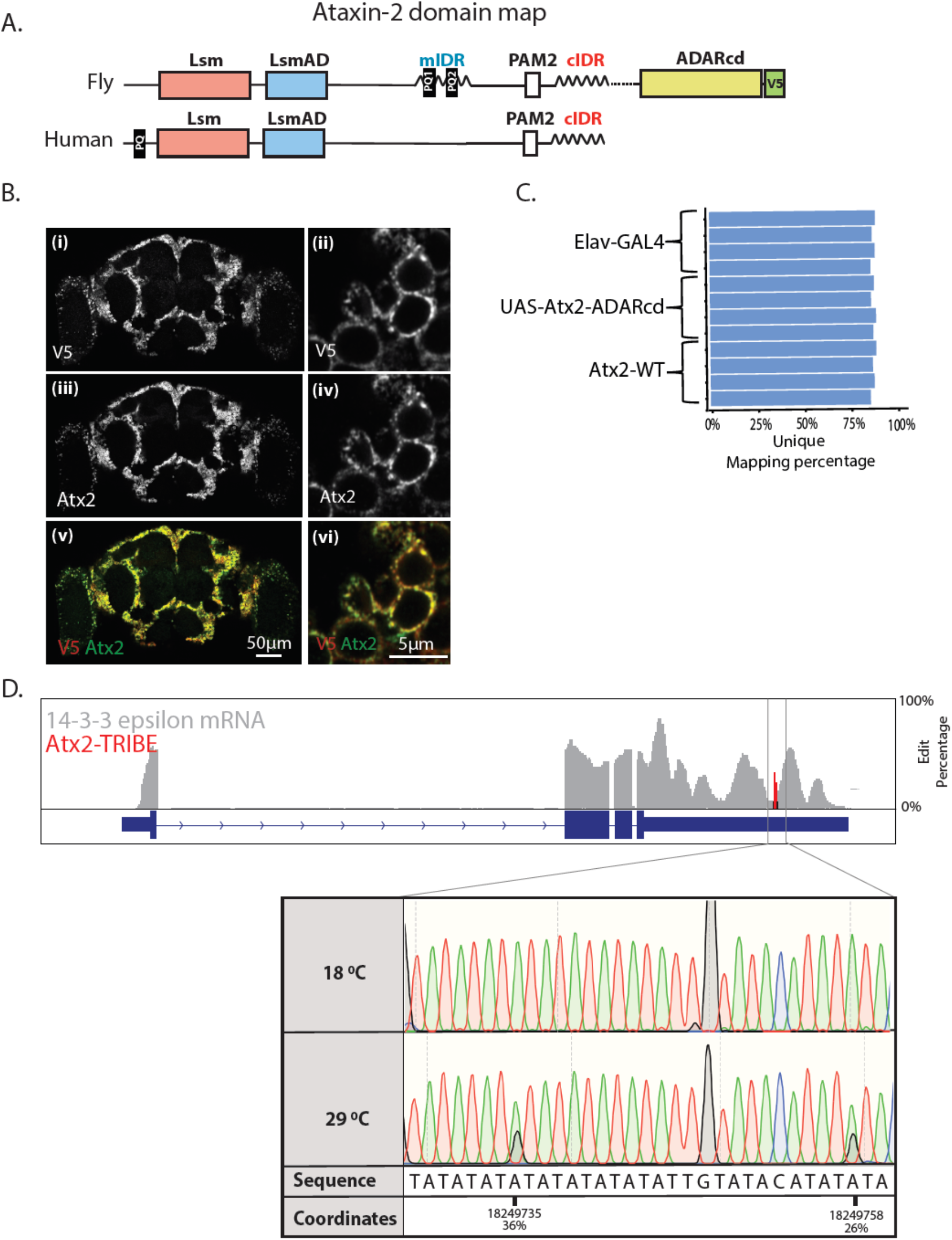
Atx2-ADARcd fusion protein expression and RNA sequencing. (A) Schematic showing the Atx2-ADARcd construct. (B) UAS-Atx2-ADARcd expression was driven using *elav-Gal4;tub-TubGal80^ts^* in adult fly brain for 5 days. Brains dissected from flies maintained at 29°C for 5 days were stained using anti-V5 and anti-Atx2 antibodies. (C) Total RNA was isolated from 10-12 brains for each replicate. Poly(A)-enriched mRNA was sequenced using paired-end libraries on Illumina 2500. The reads obtained were of high quality and mapping to *Drosophila melanogaster* genome was => 80% for all the samples. (D) An example of a gene, *14-3-3 epsilon*, showing multiple edits specifically in the exons. Red lines show the edit percentages at the different modified nucleotides with respect to the total mRNA present at a given moment of time.

**Supplementary Figure 2:**
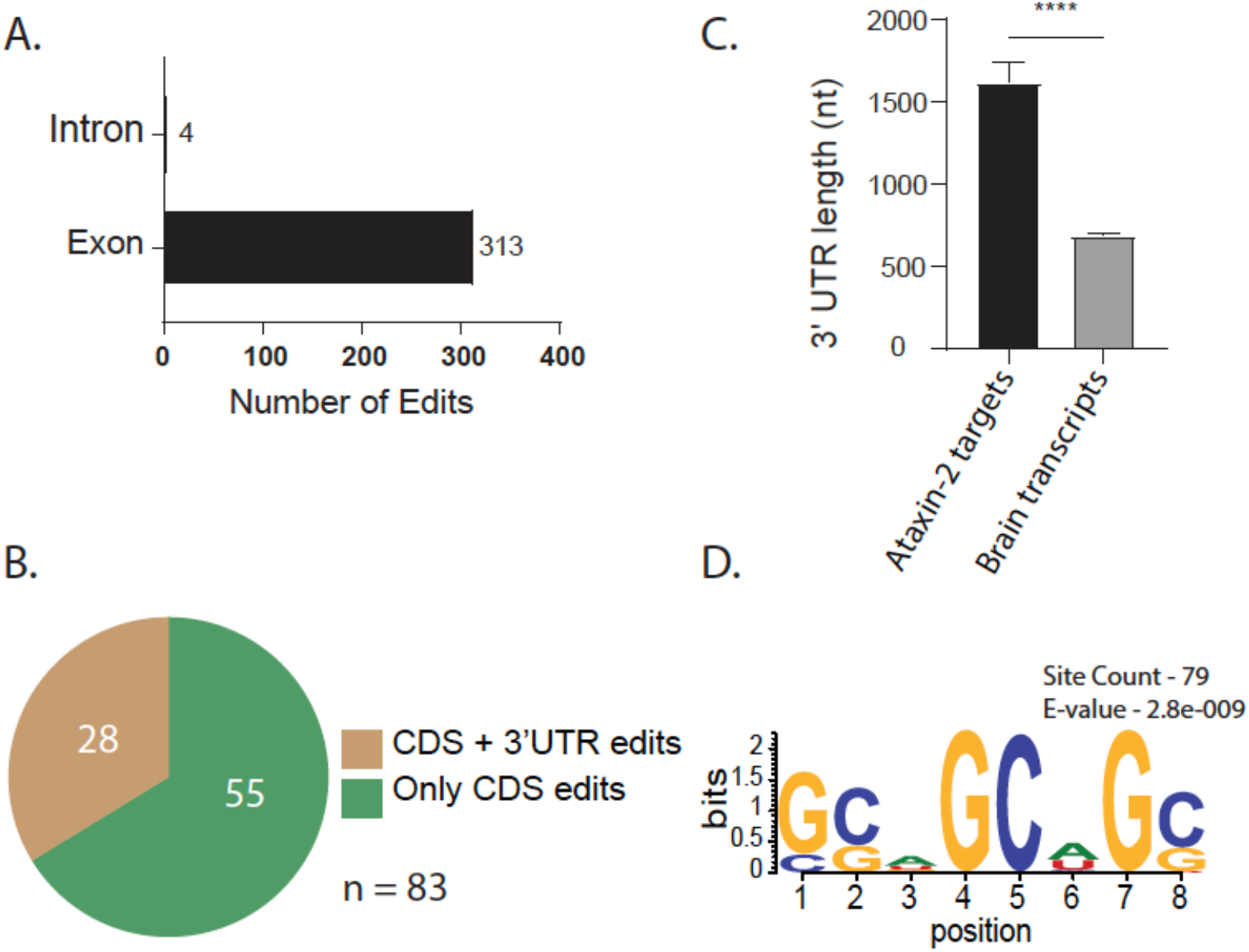
Exon-specificity of Atx-2 mRNA interactions and sequence selectivity of observed Atx-2-coding sequence (CDS) interactions. (A) Atx2-ADARcd fusion protein exclusively edits exons indicating cytoplasmic binding to mature mRNAs. (B) Coding sequence edits often occurred together with a 3’UTR edit on the same mRNA consistent with an initial Atx2-3’UTR interaction enabling edits to a proximal coding sequence as well. (C) Atx2-target mRNAs have longer than average 3’UTRs. Mann-Whitney U test was used to determine the difference in the 3’UTR length between Atx2 targets and brain transcriptome. p-value is <0.0001 and error bars represent SEM (D) A GC-rich motif for Atx2-CDS association indicated by analyses of sequences flanking CDS edit sites.

**Supplementary Figure 3:**
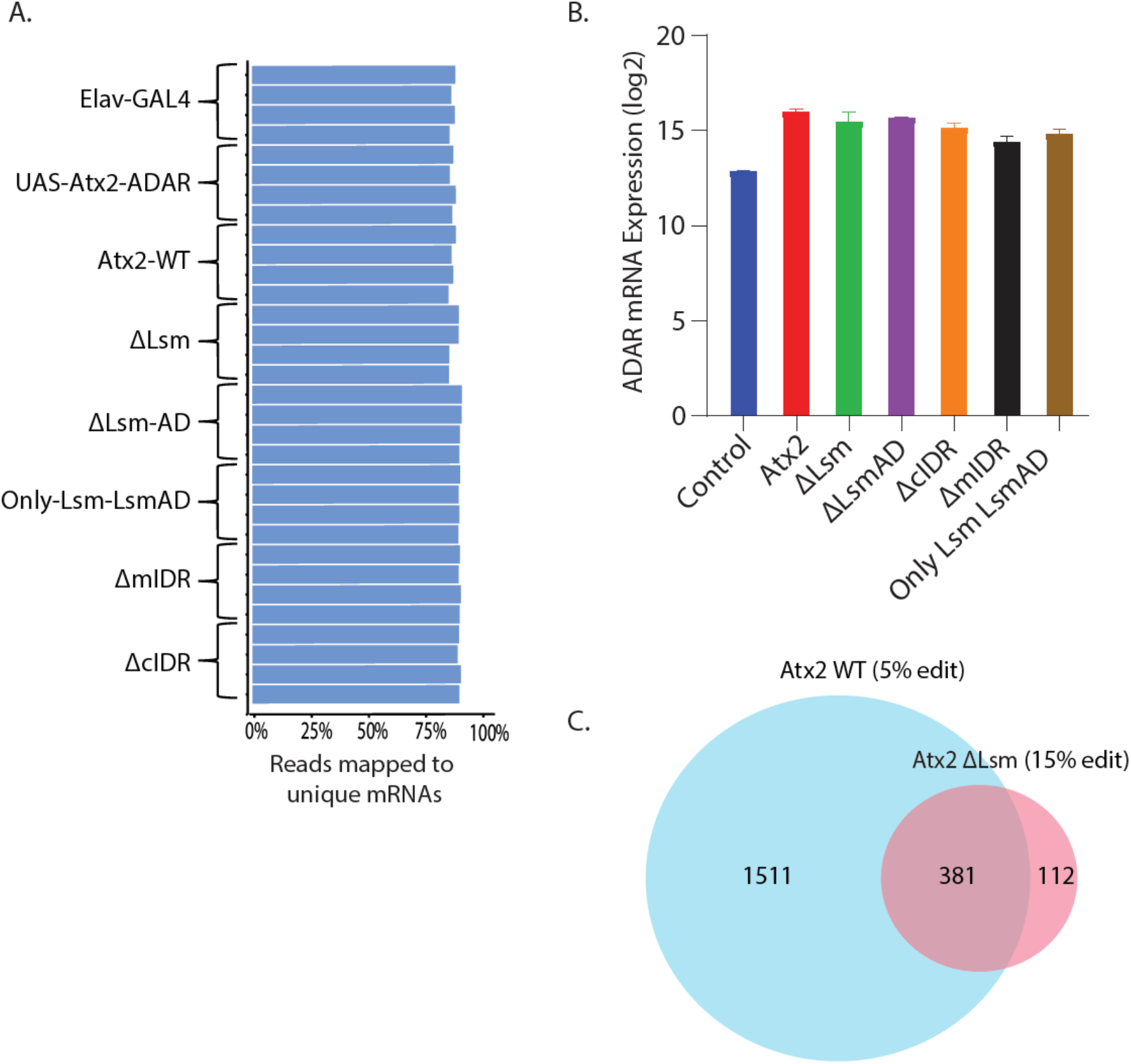
Equivalent read quality and ADAR expression across Atx2 deletion analyses. (A) Poly(A) enriched mRNA from 10-12 brains was sequenced using paired-end libraries on Illumina 2500. The reads obtained were comparable between genotypes and of high quality and mapping to the *Drosophila* genome was => 80% for all the samples (Supp. Table 5). (B) Quantification for ADAR mRNA as a read out for transgene expression showed expression above the endogenous control and comparable between Atx2 WT and the various mutant transgenic lines. (C) Atx2ΔLsm unique targets were present at reduced edit threshold in Atx2 WT. Several of the Atx2ΔLsm edits found at 15% threshold was found in Atx2 WT edits at 5% threshold.

**Supplementary Figure 4:**
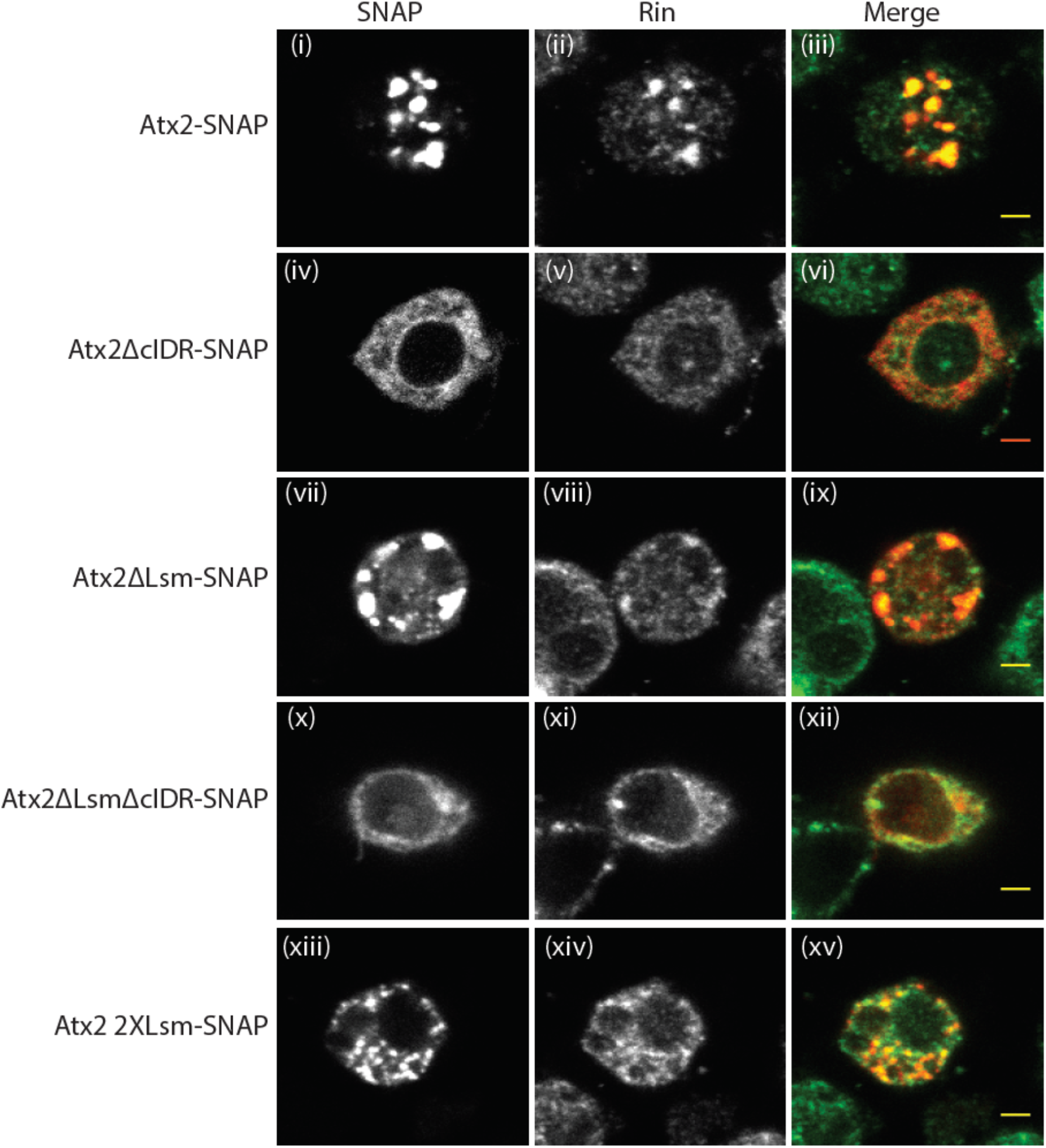
Granules formed by over-expression of Atx2 and ΔLsm resemble SGs that also contain SG protein Rasputin (Rin). Atx2WT forms cytoplasmic granules (i) which contain Rin (ii and iii). The deletion of cIDR blocks granules formation (iv, v and vi). Atx2 granules formed upon Lsm domain deletion also resemble Atx2WT granules and contain SG protein Rin (vii, viii and ix). The deletion of both Lsm and cIDR domains block the ability of Atx2 to form granules (x, xi and xii). Atx2-2XLsm where mIDR was replaced with a second Lsm domain reduces Atx2 granule size, but resemble WT granules and contain Rin (xiii, xiv and xv). The scale bars correspond to 2µm.

